# Growing a Root from Its Genome: Emergent Tissue Layers and Zones in a GPU-Native Simulation of the *Arabidopsis* Root Tip

**DOI:** 10.64898/2026.11.05.764577

**Authors:** Emeline Auger, Mickaël Gandecki, François-Xavier Heng

## Abstract

Models of the *Arabidopsis* root tip usually take its cell identities as given. Here we grow a root tip from a single genome. In a two-dimensional median section, every cell reads the same network of 62 genes (26 core genes from an earlier, unpublished model of callus and 36 added for the root) and starts with the same expression; the only spatial cues are the polarity of auxin transport, auxin from the shoot and the free surface. Identities and zones are never given to the simulation; they are read from gene expression. Cells are polygons of a shared mesh, relaxed towards mechanical equilibrium under turgor at every step. In 48 simulated hours, a tip-shaped template grows into a root of 3 000–4 500 cells with epidermis, cortex, endodermis, pericycle and stele in order along most of its length, SHORT-ROOT and SCARECROW in the endodermis, an auxin maximum, a quiescent centre and a cap at its tip, and meristem, elongation and mature zones. A WOX5 gate built from published interactions and one hypothesis keeps a quiescent centre to 48 h in 17 of 20 simulated roots that were not used for tuning (85%; Wilson 95% interval 64–95%; criteria fixed before the run), against 9 of 20 roots used for tuning, before the gate. The niche is not maintained beyond 48 h: grown on to 73 h, only 6 of these 20 roots still have a quiescent centre (30%; 15–52%). All pre-registered criteria hold in only 1 of 20 roots, because the quiescent centre is too wide at 0 h; without that count, 12 of 20. The meristem is too long, and cortical cells made after the start of growth elongate only three- to four-fold instead of eight- to thirty-fold, a limit set in large part by the numerical growth scheme. Mutants are a major weakness: several look like their published phenotype, but an exploratory re-run of a mutant benchmark fixed in advance for our first model passes one of nine predictions. Nor does the root regenerate: cut near its tip, it re-forms an auxin maximum and a columella but no quiescent centre. On a first, simpler model and without any polarity cue, a mass of identical cells still sorts into concentric layers and, when PIN carriers follow the auxin flux, concentrates auxin into a few foci, but forms no stem-cell niche: the genome lacks a lateral inhibition between foci. We state the limitations and the next stages.

## 1 Introduction

The *Arabidopsis* root tip is one of the best described organs in developmental biology. Its concentric layers, its stem-cell niche and its zones of division, elongation and maturation have each been tied to a few transcription factors, hormones and their movements between cells [Dolan et al., 1993, Helariutta et al., 2000, Sabatini et al., 1999, Aida et al., 2004, Sarkar et al., 2007, Dello Ioio et al., 2007]. Computational models have reproduced many of these mechanisms, but most fix cell identities: cells are told what they are, and the model computes what happens next [Grieneisen et al., 2007, Mähönen et al., 2014, Cruz-Ramírez et al., 2012]; others self-organise one feature, such as the auxin maximum or the zonation, on a given tissue [Mironova et al., 2010, Salvi et al., 2020]. Here we ask the opposite question: what does a root look like if no cell is ever told what it is?

We grow a two-dimensional median section of the root tip in which every cell reads the same gene network of 62 genes and hormone pathways and starts with the same expression. Identities and zones are never given to the simulation; they are read from each cell’s expression and contacts. Mutants are made only by editing the genome. The simulator is written for GPUs, so that many roots can be grown side by side and compared with quantitative targets.

This work continues the approach of our first preprint [Auger et al., 2026], in which one genome read by every cell drove a simulated root, but it uses a new simulator and a new network. The number of genes is not carried over from that work, whose network had 35 genes: the 62 genes here are 26 core genes (auxin transport and response, lateral-root initiation, PLETHORA, WOX5, SHORT-ROOT, SCARECROW, the epidermal regulators WER and GL2, and the cell cycle), first written for an earlier, unpublished model of undifferentiated callus, and 36 genes added for the root (Section 2). That callus model is distinct both from our first preprint and from the “first model” of Sections 3.4 and 3.5, an earlier version of the present simulator in which cells are bonded centres.

We report what emerges, how it compares with the real root, how robust it is on replicate simulated roots set aside in advance and never used for tuning (Section 2), and how far it is from published mutant phenotypes.

## 2 A Root Built from One Genome

The simulation represents a median longitudinal section of the root tip. Its inputs are deliberately few (Figure 1):

- **One genome**, identical in every cell: 62 genes, each with a promoter that reads transcription factors and signals, mRNA and protein half-lives, and the cellular behaviour its product drives (division licence, wall loosening, maturation, cell death, auxin transport). Seven products move between cells, among them SHORT-ROOT (SHR), WOX5 and CLE40. The genes are grouped in 17 modules. Of the 62, 26 core genes come from an earlier, unpublished model of callus (Introduction), and the 36 added for the root cover, by function, the niche and the cap (PLT2, PLT3, CLE40, ACR4, FEZ, SOMBRERO, BRN), cytokinin and its crosstalk with auxin (IPT, CKX, ARR1, ARR12, SHY2, GH3.17, AHP6, ARR5), radial patterning (PHB, miR165, JKD,
- ATML1, LHW, CYCD6), auxin synthesis and PIN polarity (TAA1, YUCCA, PINOID, WAG, PP2A, D6PK, SOSEKI), growth and maturation (EXPA1, BZR1, RGA, a maturation gene, UPB1), the cytoskeleton (KTN1, CLASP) and the DR5 reporter. All 62 are active in the vertex-model runs reported here; in the runs of the first model (Section 3.4) the maturation regulator UPB1 is silent.
- **Cell physics** (a vertex model): cells are polygons of a shared mesh, relaxed towards mechanical equilibrium under turgor at every step (Methods); walls yield only in tension and above a threshold [Lockhart, 1965], and cortical microtubules orient growth. Auxin moves by diffusion and by PIN-mediated efflux through walls.
- **A starting tissue in which every cell has the same gene expression**: a template of a root tip of 226–228 cells. Before growth, the network runs for 72 h on this fixed geometry, so identities at 0 h are already an output of the genome.
- **Three spatial cues**: the tipward polarity of auxin efflux, which cells inherit and which also orients their microtubules; auxin supplied at the shoot end; and the free surface of the organ.

**Figure 1.**
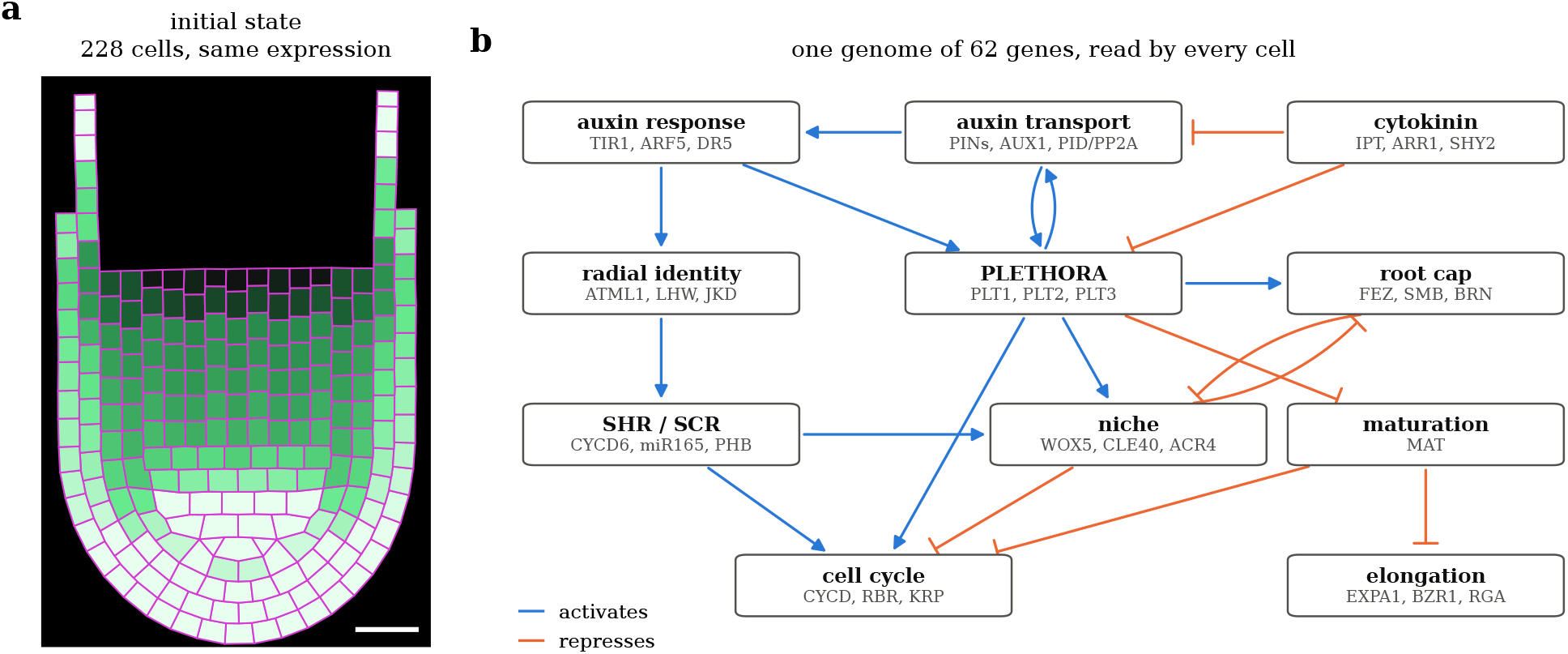
The inputs. (a) The starting template after the network has run for 72 h on its fixed geometry; every cell started with the same expression (walls in magenta, auxin response DR5 in green; scale bar 20 µm). (b) The main interactions between gene modules, read by every cell; arrows activate, bars repress. The grown root is shown in Figure 2.

Everything else is forbidden. The simulation contains no tissue label, no zone and no rule of the form “if this is a stele cell, then”. A real radicle is already patterned at germination [Scheres et al., 1994]; our template is not, and its patterning is part of what the network has to do.

### Identities and zones are read-outs

To measure the root, each cell is labelled from its own expression and contacts with fixed thresholds: epidermis (WER or GL2), cortex (JKD without SCARECROW, SCR), endodermis (SCR next to the stele), stele (SHR promoter or PHB, without SCR), pericycle (stele cell touching the endodermis), lateral root cap and columella (SOMBRERO), quiescent centre (WOX5 with a WOX5-expressing neighbour) and initials (neighbours of the quiescent centre). Zones are read from the division licence and the maturation lock. These labels are never read back by the simulation; we checked that exchanging them leaves the simulated root unchanged.

### Replicate roots and seeds

The simulation is stochastic: the positions of the template cells are jittered (by up to 8%), and divisions carry noise. Each run draws these from a random-number generator started from an integer, its *seed*. A seed is therefore only the label of one simulated root, not a plant seed: two seeds give two different roots from the same genome and the same settings, as two plants of one genotype differ, and the same seed gives the same root again when it is run on the same machine with the same number of roots grown together; on another machine the root can differ, sometimes enough to change a result (Methods). Seeds are numbered in sets. Seeds 1–4 and 101–120 (the development set) were used to develop and tune the model. Seeds 121–140 (the held-out set) were set aside before the final settings and never used for tuning; they were run once for the validation of Section 3.2, and no setting was changed afterwards. The same seeds were then grown again, with the same settings, for the figures, the mutants and the longer runs and regeneration test of Section 4.

## 3 Results

The roots shown are the simulated roots of seeds 121, 122 and 123. A seed is only the number of the random draw that makes one replicate root (Section 2); these three are the first three of the held-out set of Section 3.2, taken in order and not selected for their appearance.

### 3.1 A layered root on wall mechanics

Figure 2 follows one root from 0 to 48 h. The three roots grow from 226–228 cells to 699–701 cells at 12 h, 1 734–1 905 at 24 h, 2 706–3 456 at 36 h and 3 255–4 472 at 48 h. At 0 h the identities are already an output of the network on the template geometry: epidermis, a central stele, a few endodermis and cortex cells, and at the tip columella (16–18 cells), initials (18–31) and a quiescent centre that is far too wide (19 cells in each root, against 2–4 in a real median section, four in all; Dolan et al. 1993). By 24 h every layer is present along the root. At 48 h the roots hold 1 302–1 865 stele, 312–407 pericycle, 338–459 endodermis, 507–774 cortex and 637–754 epidermis cells, with 10–12 columella and 5–10 lateral root cap cells at the tip. The quiescent centre has narrowed to 2 and 6 cells in two roots and is absent in the third (seed 122) at 48.2 h, although that seed kept one in the validation run (Methods). In the root of Figure 2, cortex fills the centre of the tip below the stele from 36 h: the layers are in order along most of the root but not in its distal 200 µm.

**Figure 2.**
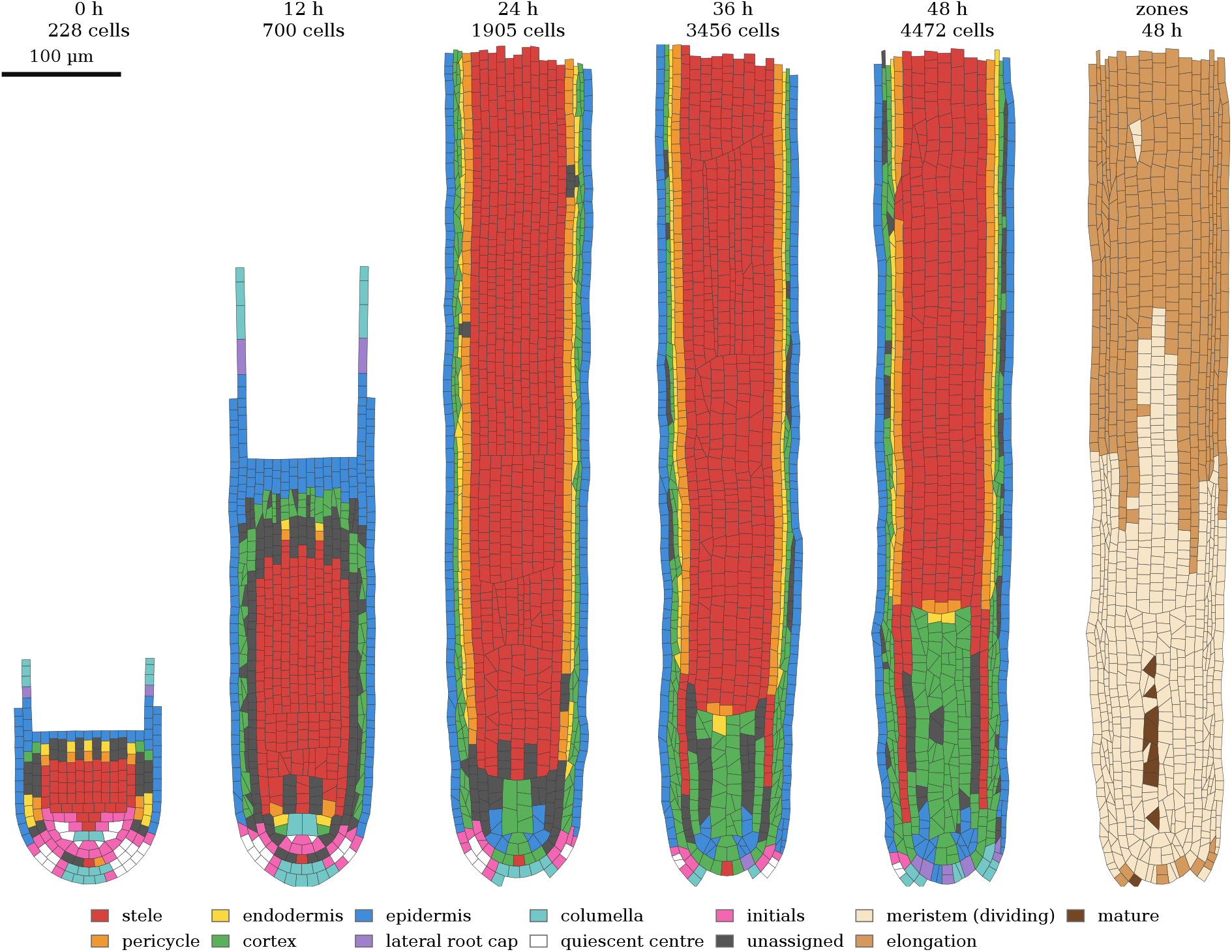
A root grows on wall mechanics under turgor. The distal 700 µm of one root (seed 121) every 12 h from 0 to 48 h, tip down, same scale; last column: zones read at 48 h. Cells coloured by identity read-out (Section 2); unassigned cells (dark grey) express no complete marker combination.

Figure 3 shows the gene products behind this pattern in the tip of the same root at 48 h (median protein levels per identity, arbitrary units). PLT1 is high in the quiescent centre, initials and cap (1.09–1.26) and low in the layers (0.04–0.15). WOX5 is confined to the quiescent centre (0.51, against (*≤* 0.08 elsewhere). The stele programme LHW is on in the stele and pericycle (0.35) and off elsewhere ≤ 0.004). SHR is made in the stele (0.16) and accumulates in the next layer (0.75), where SCR is on (0.90, against ≤ 0.03 in every other layer). The ground-tissue programme JKD marks endodermis and cortex (0.90 and 0.47), and ATML1, a marker of the embryonic protoderm in the plant [Lu et al., 1996], the surface and the cap (0.94–1.27, against ≤ 0.07 in the inner layers). The auxin response DR5 peaks at the tip (1.19–1.35, against 0.48–0.57 in the layers). Because the identity read-out is computed from some of these products, these domains do not validate the read-out; what they show is that the genome, read in the same way by every cell, partitions the tissue into concentric expression domains, two of which (ATML1 and SCR) are one cell layer thick, as the protoderm and the endodermis are in the plant [Lu et al., 1996, Helariutta et al., 2000, Di Laurenzio et al., 1996, Nakajima et al., 2001].

**Figure 3.**
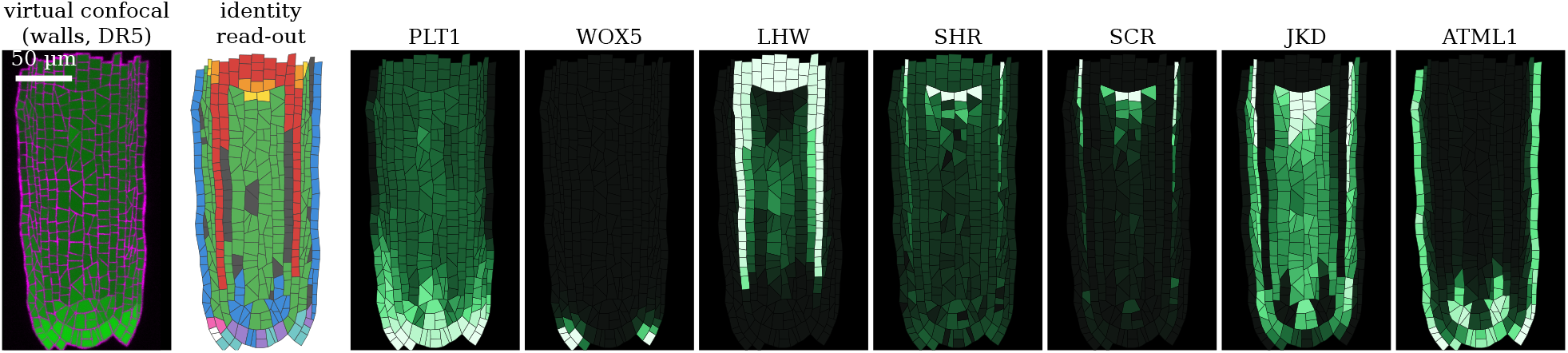
Expression domains in the tip. The distal 260 µm of the root of Figure 2 at 48 h, tip down. From left: virtual confocal section (walls in magenta, DR5 auxin response in green; scale bar 50 µm), identity read-out (colours as in Figure 2), and the protein level of seven genes, each scaled between its 1st and 99^th^ percentile in the section.

Figure 4a compares the roots with the target root. They form a distal meristem of dividing cells, an elongation zone and, from 36 h, a mature zone (Figure 2, last column, at 48 h). The meristem is too long: the first cortical cell twice as long as dividing cortical cells lies 743–896 µm from the tip (Methods), against 180–350 µm at about five days, measured from the quiescent centre to the first elongated cortical cell [Dello Ioio et al., 2007, Perilli and Sabatini, 2010]. Cells leave the cycle earlier, 250–500 µm from the tip, but cortical cells only double their length 240–570 µm further up: the transition between division and elongation is too long.

**Figure 4.**
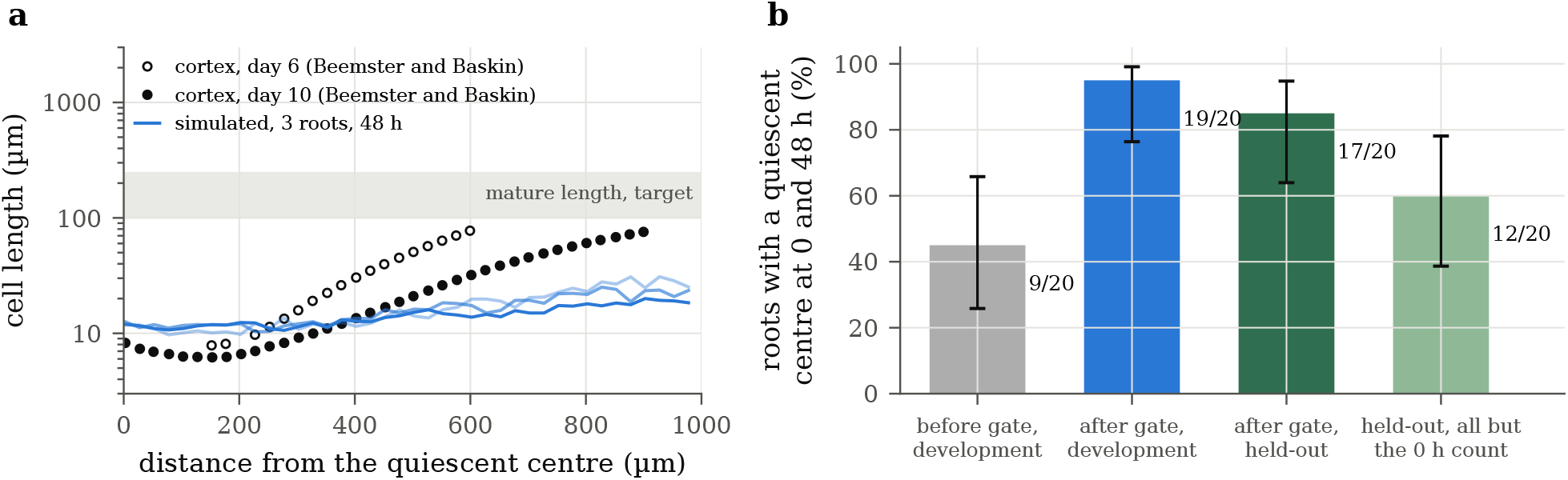
Cell length and niche maintenance. (a) Median length of all cells (all files and the root cap; the cortex alone differs by at most 5–8 µm, see text) against distance from the quiescent centre in the three simulated roots at 48 h (distance from the tip minus 10 µm, the depth of the quiescent centre in the two roots that have one at 48 h; the same offset is used for the third), and cortical cell length in real roots six and ten days after germination, digitised from Figure 3A of Beemster and Baskin [1998]; the grey band is the target for mature cells. (b) Roots keeping at least one quiescent-centre cell at 0 and 47.6 h, before and after the WOX5 gate, on the development roots (seeds 101–120) and on the held-out roots (seeds 121–140) run once; last bar: all pre-registered criteria except the quiescent-centre count at 0 h, which fails on 18 of the 20 held-out roots. Error bars: Wilson 95% intervals.

Cells also elongate too little. At 48 h the median mature cortical cell is 60–62 µm long (63–70 µm at 36 h; target 100–250 µm), but this median mixes two cohorts. Cortical cells born before 24 h, from the starting template, left the cycle almost together and are 75–85 µm long; cortical cells born after 24 h, the first made by the established meristem, are 41–46 µm long, 3.2–4.1 times the length of dividing cortical cells (target 8–30). At 36 h every mature cell still belongs to the first cohort, so the fall of the median between 36 and 48 h reflects the arrival of the second cohort, not a change in how cells grow. The length of the second cohort is set in large part by the numerical scheme: a wall extends its rest length by at most its elastic strain (about 3%) per step of 0.28 h, which caps its relative growth rate near 0.11 h^*−*1^, and growth stops 16 h after a cell starts to mature (Methods). We have not tested how this cap changes with the time step, so this limit should not be read as a property of the genome.

Close to the quiescent centre, simulated cells are about as short as real ones (10–12 against 6–8 µm), but their median length then rises slowly and stays at or below about 30 µm within 1 mm. This profile pools all cell files and the cap; the cortex alone gives nearly the same profile (within 5 µm, and 8 µm in bands holding only a few cortical cells). In real roots, cortical cells reach 40 µm about 450 µm from the quiescent centre at six days and 700 µm at ten days (Figure 4a), and keep elongating to the end of a growth zone 1.2–2.3 mm long [Beemster and Baskin, 1998].

These comparisons with roots five to ten days old are made during the start-up of growth: the simulated roots double in length between 36 and 48 h (from 4.2–4.5 to 8.2–10.0 mm), and 54–58% of their mature cells come from the starting template, whose cells left the cycle almost together. Only cells born after 24 h can be compared with an established root.

### 3.2 A WOX5 gate keeps the niche

On the twenty development roots (seeds 101–120, used for tuning), the vertex model before the WOX5 gate kept a quiescent centre at both 0 and 48 h in only 9 roots (45%; Wilson 95% interval 26–66%). In the others it was absent from the start (6 roots) or turned into cap. The cause was in the genome: the WOX5 promoter ran at about 8% of its maximal rate in the niche, and its protein (0.23) sat just above the read-out threshold (0.2), with nothing separating niche from endodermis or columella except distance to the auxin maximum. A jitter of 8% in initial positions was enough to tip it. We changed only gene-network parameters, most following the literature: WOX5 requires PLT [Aida et al., 2004, Ding and Friml, 2010] and now reads the PLT3 dose cooperatively, after the dose-dependent action of PLT proteins [Galinha et al., 2007] (the cooperativity and the choice of PLT3 are model choices); auxin represses WOX5 directly, standing for its repression through ARF10 and ARF16 in the plant [Ding and Friml, 2010], while inducing PLT [Aida et al., 2004]; CLE40 from the columella represses WOX5 more strongly [Stahl et al., 2009]; WOX5 represses the cap regulator SMB [Bennett et al., 2014] and, as a hypothesis, SMB represses WOX5, which makes the niche–columella boundary bistable; and WOX5, which is mobile [Pi et al., 2015], is imported into the nucleus faster, by analogy with the nuclear retention of SHR [Cui et al., 2007], which reduces its spread. No position rule, identity or read-out threshold was changed.

With success criteria written in advance (Methods), we then grew twenty new roots, from seeds 121–140 that had never been used during development or tuning, once each, with the final settings. With the gate, a quiescent centre was present at 0 and 48 h in 19 of the 20 development roots and in 17 of the 20 held-out roots (85%; 64–95%; Figure 4b). The development roots measure how well the gate was tuned; only the held-out roots test it. Twenty roots per set give wide intervals (Methods). All pre-registered criteria held in only 1 of 20 held-out roots, because the quiescent centre is too wide at 0 h (below); without that count, they held in 12 of 20 (60%; 39–78%). Of the three roots without a quiescent centre, one (seed 137) is a numerical failure: the computation broke down at the start of growth, the root never grew, and the same failure occurs without the gate; the other two lose the quiescent centre by 48 h. These counts hold at 48 h only: grown on to 73 h, the quiescent centre is lost in most of these roots (Section 4). As controls fixed in the same protocol, *shr* and *scr* genomes (SHR or SCR dose set to zero) were grown with the gate on seed 1: neither has any cell read as endodermis at 0 or 48 h, so the gate does not restore the endodermis without them. One limitation remains: at 0 h the quiescent centre read-out covers the whole bottom row of the template (10–38 cells instead of 2–4), because the PLT3 plateau of the model is nine files wide. A steeper gate that kept only the central cells switched off later, as the quiescent cells grew and diluted WOX5.

### 3.3 Mutants made by editing the genome

Twelve genome edits were grown on the vertex model to 48 h on one seed (121), with the same settings as the wild type; Figure 5 shows the ten that could be read. These are qualitative observations, not statistical tests: for this figure each mutant was grown once and compared with one wild-type root, without criteria fixed in advance, so a difference may hold for that root only. A phenotype such as the loss of the endodermis is described as seen in that root; for *shr* and *scr* it was also seen in a second root of each, the controls of Section 3.2 (seed 1). In *shr* and *scr* the simulation loses the endodermis, and with it the pericycle read-out, which requires an endodermal neighbour; in the plant the ground tissue is reduced to one layer, of cortex identity in *shr* and of mixed identity in *scr*, and the pericycle is kept [Benfey et al., 1993, Helariutta et al., 2000, Di Laurenzio et al., 1996]. In *scr* much of the inner tissue is left unassigned, and the *shr* root has more cells than the wild type (7 260 against 4 472) instead of fewer. In *wox5* the layers are kept, the quiescent centre is lost and the columella shrinks (5 cells against 12); the real mutant differs, as its columella stem cells differentiate and its quiescent-centre cells persist but divide [Sarkar et al., 2007, Forzani et al., 2014]. In the first model the same edit lost the stele. In *fez* the root has no columella or lateral root cap cell, a stronger defect than the fewer cap layers of the real mutant [Willemsen et al., 2008]. In *plt1 plt2* the template does not grow at all, a stronger defect than the real double mutant, whose embryonic root pole is already abnormal and whose meristem differentiates within days of germination [Aida et al., 2004]. Among the meristem-size mutants, *wei8 tar2* nearly loses its meristem (20 dividing cells against 583) and its root stays short (1 448 cells), as published [Stepanova et al., 2008, Brumos et al., 2018], while *arr1 arr12* and *ipt3 ipt5 ipt7*, which enlarge the real meristem [Dello Ioio et al., 2007], have fewer dividing cells than the wild type (520 and 374), and *shy2-2*, which shrinks it [Dello Ioio et al., 2008], has more (632). These meristem read-outs are taken on one seed at 48 h, during the start-up of growth, when a mutant can shift the timing of the first wave of differentiation rather than the size of the established meristem, and their direction depends on the read-out: *arr1 arr12* and *ipt3 ipt5 ipt7* have fewer dividing cells than the wild type, but their first elongated cortical cell lies farther from the tip (1 178 and 1 654 against 896 µm), because their cells elongate less. We do not interpret the direction of these three phenotypes. Several mutant roots are also further from mechanical equilibrium than the wild type at 48 h (Methods). In *ktn1* the cells swell radially, as in katanin mutants [Burk et al., 2001, Bichet et al., 2001], but the root proliferates to the simulation’s capacity of 8 192 cells, whereas the real mutant roots are short. Two edits, *brn1 brn2* and *pin1 pin3 pin7*, could not be read at 48 h on this seed: the read-outs did not find their tip on the mesh.

**Figure 5.**
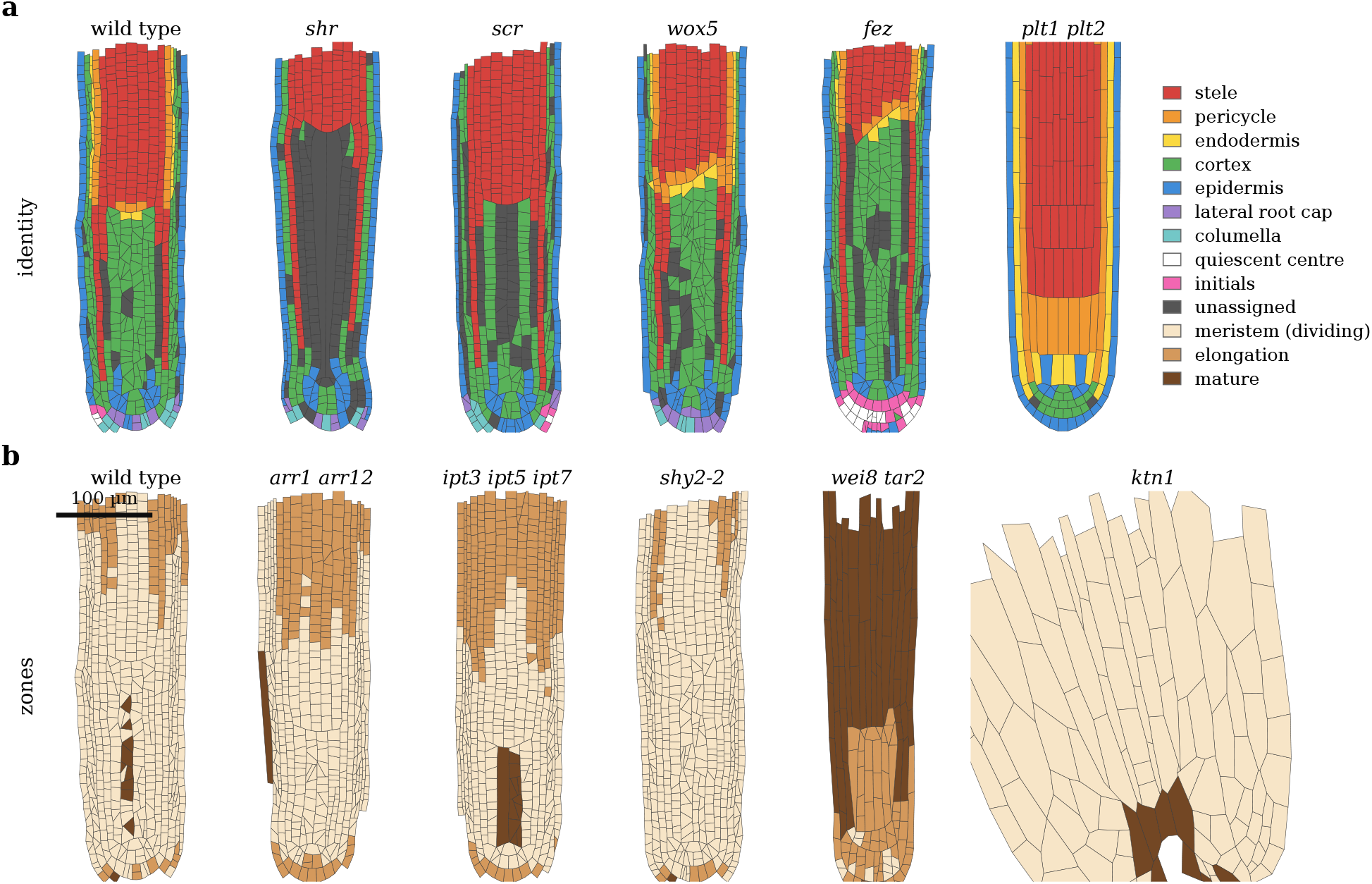
Mutants made by editing the genome, vertex model, 48 h, seed 121, distal 400 µm, tip down, one root each. (a) Radial pattern, niche and cap, coloured by identity: *shr* and *scr* lose the endodermis and pericycle, *wox5* keeps its layers and loses the quiescent centre, *fez* has no cap, *plt1 plt2* does not grow. (b) Zones: *wei8 tar2* nearly loses its meristem, as published; for *arr1 arr12, ipt3 ipt5 ipt7* and *shy2-2* the direction of the change depends on the read-out (text); *ktn1* (wider panel, same scale) swells radially. Scale bar 100 µm.

As an indication only, we re-ran on eight seeds at 48 h the nine counted predictions of a mutant benchmark that the vertex model can read out (each an interval on one read-out relative to the wild type; Table 1). The benchmark and its pass criteria were written down before any run, but for our first model; on the vertex model, which did not exist then, this re-run is exploratory (a “dry run”) and is not reported as a test of the vertex model. One prediction passes (*brn1 brn2* : more columella tiers; Bennett et al. 2010), against three of nine for the first model at 140 h. Four fail in the wrong direction or without effect for reasons of model structure: *jkd* loses ground tissue instead of gaining it [Welch et al., 2007], *lhw* loses the whole stele instead of part of it [Ohashi-Ito and Bergmann, 2007], *acr4* has no effect on columella [De Smet et al., 2008], and *pin1 pin3 pin7* enlarges the meristem instead of shrinking it [Blilou et al., 2005]. We trace these to single model genes that stand for whole gene families and to a division licence that does not read the auxin level. Generic structural edits brought the count to two of nine but reduced niche maintenance in the wild type (7 of 10 wild-type roots, seeds 101–110, against 10 of 10 without the edits), so we did not adopt them. Mutant phenotypes are now the weakest part of the model.

**Table 1.** Counted mutant predictions of the pre-registered benchmark. Estimate: ratio or difference to the wild type on the predicted read-out, with its 95% interval. A prediction passes when the estimate lies in the pre-registered interval and its interval excludes no change. *†*Failed because the mutant was rejected for integrity more often than the wild type. The vertex-model run is exploratory (a dry run) and does not count.

|  |  | first model (32 seeds, 140 h) |  | vertex model, dry run (8 seeds, 48 h) |  |
|---|---|---|---|---|---|
| edit | predicted interval | estimate [95% CI] | pass | estimate [95% CI] | pass |
| P01 *jkd* | [0.15, 1.50] | −1.79 [−2.37, −1.28] | × | −0.93 [−3.03, −0.54] | × |
| P02 RBR reduced | [1.10, 3.00] | 1.54 [1.50, 1.61] | ✓ | 1.24 [0.88, 1.74] | × |
| P03 *lhw* | [0.30, 0.90] | 0.00 [0.00, 0.00] | × | 0.00 [0.00, 0.00] | × |
| P04 *acr4* | [1.10, 2.50] | 1.07 [0.77, 1.24] | × | 1.06 [0.72, 1.48] | × |
| P05 *brn1 brn2* | [1.25, 6.00] | 4.18 [3.21, 5.33] | ✓ | 1.71 [1.14, 2.43] | ✓ |
| P06 polar transport inhibited | [5.00, 150] | 733 [292, 1730] | × | no read-out | × |
| P07 35S::*PID* | [0.00, 0.60] | 0.45 [0.22, 0.83] | ✓ | 0.24 [0.10, 0.32] | ×<sup>†</sup> |
| P08 *pin1 pin3 pin7* | [0.30, 0.90] | 0.41 [0.17, 0.54] | ×<sup>†</sup> | 1.45 [1.29, 1.74] | × |
| P09 *arr12* | [1.05, 1.60] | 0.95 [0.84, 1.04] | × | 0.96 [0.74, 1.28] | × |
| passes |  |  | 3/9 |  | 1/9 |

### 3.4 A mass of cells without an axis

The root above inherits one cue from its starting tissue: the tipward polarity of auxin efflux. To ask what the genome does without it, we grew, on the first model (Section 3.5), a block of 288 cells with the same expression and no polarity at all (PIN spread evenly over all faces; Figure 6). Without added hormone the mass proliferates to more than 10 000 cells in 72 h and still sorts itself into concentric layers: epidermis outside, a cortex, and a core of stele with patches of endodermis and pericycle (Figure 6a). This only loosely recalls real callus, which has a root-meristem-like identity and arises from pericycle-like cells [Atta et al., 2009, Sugimoto et al., 2010]. Bathed in auxin and cytokinin through its whole surface, as an explant on callus-inducing medium [Valvekens et al., 1988], it forms no stem-cell niche either. We then let cells reorient their PIN carriers along the auxin flux (canalisation, Mitchison 1981), with 24 h on callus-inducing medium followed by medium without hormone (Figure 6b–d). At 24 h, still on callus medium, the WOX5 expression that all cells carry at the start has retracted to the centre into a complete, concentric niche: quiescent centre, a ring of initials and columella (Figure 6b). This niche is a transient remnant rather than a new one: its quiescent centre and initials are lost between 36 and 48 h whatever the medium (at most one and four cells are left at 48 h, in runs kept on callus medium throughout or moved to hormone-free medium at 24, 48 or 72 h), while some columella remains. The mass keeps growing and concentrates its auxin response: at 96 h the run shown has one focus of 26 cells (Figure 6c,d). Runs kept on callus medium for 48 h or longer form larger but transient foci, of 55–115 cells at 84 h, which shrink to 5–23 cells by 96 h and are gone by 108 h. WOX5 is never induced again on these foci. Reorienting PIN towards neighbours richer in auxin instead [Jönsson et al., 2006, Smith et al., 2006], with the same 24 h on callus medium, breaks auxin into dozens to over a hundred local maxima (123–130 in the run shown and its replicate), with a dozen or more small WOX5-expressing groups of one or two cells, and arrests growth at about 3 000 cells (Figure 6e). Neither the WOX5 gate of Section 3.2, whose threshold the weak callus foci never reach, nor a stronger auxin-synthesis loop, which runs away through the whole tissue because PLETHORA is high in every cell, regenerated a niche. What the genome lacks is a lateral inhibition by which one focus strengthens itself at the expense of its neighbours.

**Figure 6.**
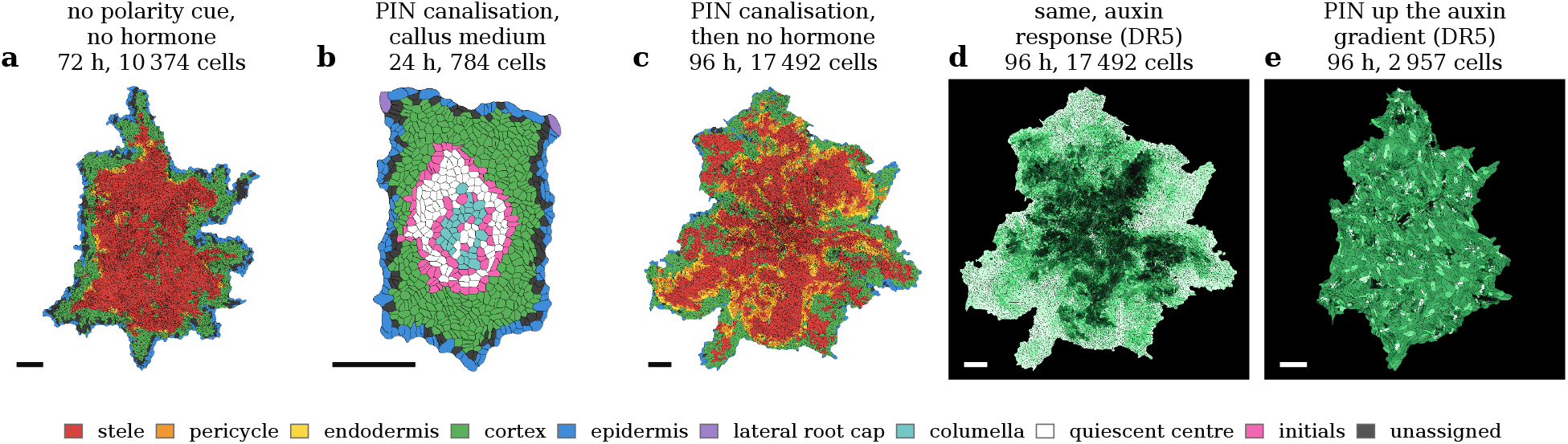
A mass of identical cells without any polarity cue. 288 cells with the same expression, PIN spread evenly over all faces (first model; colours as in Figure 2; scale bars 100 µm). (a) Without hormone, 72 h: concentric epidermis, cortex and stele. (b–d) With PIN canalisation, 24 h on callus-inducing medium then medium without hormone: a transient central niche at 24 h, at the end of the callus medium (b), and at 96 h layered tissue (c) and its auxin response (d, DR5), uneven across the mass, with one focus of 26 cells. (e) With PIN pointing up the auxin gradient, same media, 96 h: auxin response broken into more than a hundred local maxima, growth arrested.

### 3.5 A first model, and what it taught us

Before the vertex model, we grew the same kind of genome on a simpler mechanics, in which cells are bonded centres, from an embryo of 31 identical cells. It formed the same five layers, an auxin maximum, a transient niche and three zones, but its cells slid past each other, its niche was lost by 140 h, and its cells elongated only two-to four-fold. Calibration against thirteen quantitative targets, with about 16 000 roots grown in two GPU campaigns, found no parameter set that reached every target, and every route to longer cells that we tried (faster wall loosening, later maturation, more stable identity proteins) cost radial order. These results motivated the vertex model, which keeps the layers; its cells elongate more when cells of the starting template are included, but cortical cells made after the start still elongate only three-to four-fold (Section 3.1).

## 4 Discussion

### What one genome builds

From cells with the same expression and three spatial cues, 62 genes read by every cell produce the layers, the niche, the cap and the three zones of the root tip, without any rule about a cell type. The first, bonded-centre model gave the same layers (Section 3.5), so the radial pattern does not depend only on the vertex mechanics.

### Robustness has to be shown on held-out roots

Before the WOX5 gate the niche was held by a promoter just above the read-out threshold and was lost in more than half of the development roots; only replicate roots and criteria fixed in advance revealed this fragility. Growing the same held-out roots further, in a later test, bounds the result in time: the gate keeps the niche to 48 h, not beyond.

### Limitations

- **Cells elongate too little**. Cortical cells made after the start reach 41–46 µm, three to four times the length of dividing cells, against 100–250 µm and eight-to thirty-fold in the plant (cells of the starting template reach 75–85 µm). The growth rate of a wall is capped by the numerical scheme at one elastic extension per step, growth stops after a fixed period, and the in-plane wall formulation slows the axial growth of long cells (Methods). The meristem is too long (743–896 µm against 180–350).
- **The niche is imperfect**. The quiescent centre is too wide at 0 h (10–38 cells against 2–4), it is absent at 48 h in 3 of 20 held-out roots (lost in two, a numerical failure in one), the cap holds fewer columella tiers than the plant, and in the distal 200 µm cortex can fill the centre below the stele. Above all, the niche is not maintained beyond 48 h: grown on to 73 h (with a larger solver budget, which leaves the count at 48 h unchanged), only 6 of the 20 held-out roots still have a quiescent centre (30%; 15–52%), against 17 of 20 at 48 h, and four of these six keep a single quiescent-centre cell. We do not yet know why.
- **Mutants are a major weakness**. Several edits look like the published mutants, but others lose tissues they should keep (*jkd, lhw*) or enlarge the meristem where the real mutant shrinks it (*pin1 pin3 pin7*); for three meristem-size mutants the direction of the change depends on the read-out; and an exploratory re-run of a mutant benchmark fixed in advance for our first model passes one of nine predictions. Some model genes stand for whole gene families, and the division licence does not read the auxin level.
- **No regeneration**. Without the inherited polarity of auxin transport, a mass of cells (first model) forms layers and auxin foci but no stem-cell niche: the genome lacks a lateral inhibition between foci. The vertex model does not regenerate a niche either. In a test fixed in advance on the held-out roots, roots were cut or ablated at 28 h and read 20 h later, at 48 h. Roots cut 130 µm from the tip re-formed an auxin maximum (18 of 20) and a columella (19 of 20), but no quiescent centre (0 of 20), whereas the plant forms a new one within about a day [Sena et al., 2009]. After ablation of the whole quiescent centre, a quiescent centre was read again in 15 of 19 roots, after a median of 2.8 h and at the same place: its neighbours were simply read as quiescent centre, rather than a new one being specified higher up as in the plant [Xu et al., 2006]. Initials in contact with a killed quiescent-centre cell had not differentiated 45 h after the ablation, unlike in the plant [van den Berg et al., 1997], but this was read at 73 h, beyond the time to which the model is valid.
- **Scope**. The simulation is a two-dimensional median section: it cannot show the rings of cortical and endodermal cells, the xylem axis or gravitropism. The model is valid to 48 h only: with the final genome and settings (and a larger solver budget, which leaves the count at 48 h unchanged), the quiescent centre is lost between 48 and 73 h in most roots, and on earlier settings the tip lost its shape between about 50 and 120 h. All read-outs at 48 h are taken during the start-up of growth (Section 3.1). Wild-type figures show one to three roots and mutants one root each, and the validation used twenty roots per set (Methods). The genome mixes established interactions with model hypotheses, and its magnitudes were set by hand and by calibration: the result shows that such a genome can suffice, not that its parameters are those of the plant.

### Roadmap

The work proceeds in stages, each closed by an objective that can be demonstrated on the simulation (Table 2).

**Table 2.** Next stages and their demonstrable objectives.

| Stage | Feature | Demonstrable objective |
|---|---|---|
| Elongation | Cell elongation on the vertex model with the layers kept, at a time step that does not change the result | Cortical cells born after the start-up of growth reach 100–250 µm, elongation ≥ 8-fold, at a relative growth rate in the elongation zone within the range measured in the plant [Beemster and Baskin, 1998]; radial order kept |
| Niche and cap | A narrow quiescent centre and a renewed cap | Quiescent centre of 2–4 cells; 4–5 columella tiers; niche kept on ≥ 90% of held-out roots |
| Gene families | Family members split where mutants require it; an auxin-dose-dependent division licence | ≥ 5 of 9 genetic predictions of the pre-registered benchmark, without loss of niche robustness |
| Self-organised niche | Lateral inhibition between auxin foci, with PIN canalisation; re-specification of a quiescent centre | A niche regenerated after excision of the tip, and in a mass of cells without any polarity cue |
| Duration | Stable growth and a maintained niche beyond 48 h | Root growth for several simulated days with the tip intact and a quiescent centre kept on ≥ 90% of held-out roots at 96 h |
| Three dimensions | Same genome and physics in 3D | Concentric rings of epidermal, cortical and endodermal cells |

## 5 Methods

### Simulator

The simulator is written in JAX [Bradbury et al., 2018]. The tissue is held in arrays of fixed capacity, and one compiled program advances it by one step of 0.28 simulated hours: gene expression, transport, growth, division, cell death, mechanics and integrity checks. Every parameter is passed as data, so the same program grows many roots side by side on one GPU. Runs reported here used NVIDIA A40 GPUs; a root of 48 h takes about one minute.

### Gene expression

Each gene’s transcription rate is a logistic function of a basal rate, of the weighted levels of its transcription factors and of environmental signals (auxin, auxin response, SHORT-ROOT received from neighbours, free surface); mRNA and protein follow first-order production and decay, and proteins are diluted by growth. Mutants are made by setting a gene’s dose to zero or by removing regulatory terms, with all other parameters unchanged.

### Mechanics

At each step the vertex positions are relaxed towards the minimum of the elastic energy of the walls against turgor (L-BFGS, at most 500 iterations, stopped earlier on stagnation over 100 iterations). The tolerance of 10^*−*3^ N m^*−*1^ was not reached: in the states of the wild-type roots saved at 36 and 48 h, the largest residual force on a vertex was 0.4–0.5 N m^*−*1^, except 1.5 *×*10^3^ N m^*−*1^ at 48 h in seed 121. Re-equilibrating that state with 4 000 iterations changed the area of 2.6% of its cells by more than 1%, but none of the read-outs reported here that we checked (mature cell length, meristem length) by more than 0.3 µm. Growing three development roots with 2 000 iterations (and crushed dead cells made passive) changed cell numbers by at most six, mature cell lengths by less than 1 µm and meristem length by at most 3 µm in two roots and 42 µm in the third. Some mutant roots are further from equilibrium at 48 h. Wall crossings are not prevented: at 48 h, 0, 4 and 0 cells had a crossed wall in the three wild-type roots, and up to 24 in the mutants.

Walls whose strain exceeds a threshold extend their rest length, by at most their current elastic extension per step (the rest length never exceeds the current length), which caps the relative growth rate of a wall near 0.11 h^*−*1^. Cortical fibres resist transverse growth. The strain along the fibres is computed from the second moments of the cell polygon. For a long cell whose fibres are slightly tilted from its transverse axis, this couples axial elongation to the stiff fibre term with a weight of about (*L/W* sin *θ*)^2^, where *L/W* is the aspect ratio of the cell and *θ* the tilt. At 48 h this weight reaches 0.13–0.26 between 0.5 and 4 mm from the tip in two of the three roots shown, where the strain of longitudinal walls falls to 2.2–2.9% for a yield threshold of 2%. The fibres therefore also slow the axial growth of long cells. A strain measured along the deformed fibre direction removes this artefact but makes the solver fail more often, and was not used here. Cells divide when their area and division licence exceed thresholds. When a cell starts to mature, under the control of its maturation genes, a timer starts; 16 h later its maturation locks irreversibly and its walls stop growing (a duration chosen by trial, not measured). Cap cells die and are sloughed under gene control.

### Validation protocol

Each replicate root is identified by its random seed (Section 2). The WOX5 gate was tuned on seeds 1–4 and 101–120 only. Success criteria (a quiescent centre at 0 and 48 h; 1–12 quiescent-centre cells at 0 and at 48 h; initials present; endodermis and cortex counts within 30% of the median of five development roots that kept their niche before the gate; a root shape checked by eye; and, as controls, no endodermis in *shr* and *scr*) were pre-registered, that is, written down and dated in our protocol before the final settings and before any run on seeds 121–140 (they were not deposited in a public registry). Seeds 121–140 were run once for this validation; one earlier run on seed 121, with a gate later abandoned, was stopped, discarded and not counted. Criteria are evaluated at step 170 (47.6 h). Intervals are Wilson 95% intervals. Each set has twenty roots, a size fixed before the runs. This is below the 32 roots per set that our own protocol requires for a final validation, so this validation is preliminary: twenty roots give wide intervals (17 of 20 is 64–95%; 12 of 20 is 39–78%). Runs are exactly reproducible only on the same hardware and with the same number of roots grown side by side; otherwise a re-run of a seed can differ: seed 122, which kept a quiescent centre at 47.6 h in the validation run, has none at 48.2 h in the re-run of the three roots shown in Figures 2–4. The mutant benchmark was written down and frozen on the first model before it was run; its re-run on the vertex model is exploratory and is not counted.

### Read-outs

Identities and zones are read as described in Section 2. Meristem length is read on the cortical file, as in Perilli and Sabatini [2010]: cortical cells are ordered by distance from the tip, and the meristem ends at the first cortical cell where the running median length of five consecutive cortical cells exceeds twice the median length of dividing cortical cells. Mature cell length is the median length of mature cortical cells, reported separately for cells born before and after 24 h; the elongation ratio is that length over the median length of dividing cortical cells. A cortical file in a median section holds few cells (1–17 per 50 µm beyond the first 250 µm in the three roots shown), so these read-outs are noisy.

### Availability

The simulator, the genome and the analyses are available from the authors on request.

